# Big data and single cell transcriptomics: implications for ontological representation

**DOI:** 10.1101/257352

**Authors:** Brian D. Aevermann, Mark Novotny, Trygve Bakken, Jeremy A. Miller, Alexander D. Diehl, David Osumi-Sutherland, Roger S. Lasken, Ed S. Lein, Richard H. Scheuermann

## Abstract

Cells are fundamental functional units of multicellular organisms, with different cell types playing distinct physiological roles in the body. The recent advent of single cell transcriptional profiling using RNA sequencing is producing “big data”, enabling the identification of novel human cell types at an unprecedented rate. In this review, we summarize recent work characterizing cell types in the human central nervous and immune systems using single cell and single nuclei RNA sequencing, and discuss the implications that these discoveries are having on the representation of cell types in the reference Cell Ontology (CL). We propose a method based on random forest machine learning for identifying sets of necessary and sufficient marker genes that can be used to assemble consistent and reproducible cell type definitions for incorporation into the CL. The representation of defined cell type classes and their relationships in the CL using this strategy will make the cell type classes findable, accessible, interoperable, and reusable (FAIR), allowing the CL to serve as a reference knowledgebase of information about the role that distinct cellular phenotypes play in human health and disease.

## Introduction

Cells are probably the most important fundamental functional units of multicellular organisms, since different cell types play different physiological roles in the body. Although every cell of an individual organism has essentially the same genome structure, different cells realize diverse functions due to differences in their *expressed* genome. In many cases, abnormalities in gene expression form the physical basis of disease dispositions. Thus, understanding and representing normal and abnormal cellular phenotypes can lead to the development of biomarkers for diagnosing disease and the identification of critical targets for therapeutic interventions.

Previous approaches used to characterize cell phenotypes have several drawbacks that limited their ability to comprehensively identify the cellular complexity of human tissues. Transcriptional profiling of bulk cell sample mixtures by microarray or RNA sequencing can simultaneously assess gene expression levels and proportions of abundant known cell types, but precludes identification of novel cell types and obscures the contributions of rare cell subsets to the gene expression patterns present in the bulk samples. Flow cytometry provides phenotype information at the single cell level, but is limited by the number of discrete markers that can be assessed, and relies on prior knowledge of marker expression patterns. The recent establishment of methods for single cell transcriptional profiling (1, 2) is revolutionizing our ability to understand complex cell mixtures, avoiding the averaging phenomenon inherent in the analysis of bulk cell mixtures and providing for an unbiased assessment of phenotypic markers within the expressed genome.

In order to compare experimental results and other information about cell types, a standard reference nomenclature that includes consistent cell type names and definitions is required. The Cell Ontology (CL) is a biomedical ontology developed to provide this standard reference nomenclature for *in vivo* cell types in humans and major model organisms (3). However, the advent of high-content single cell transcriptomics for cell type characterization has resulted in a number of challenges for their representation in the CL (discussed in Bakken 2017 (4)). In this paper, we review some of the recent discoveries that have resulted from the application of single cell transcriptomics to human samples, and propose a strategy for defining cell types within the CL based on the identification of necessary and sufficient marker genes, to support interoperable and reproducible research.

## Application to the human brain

Initial progress in neuronal cell type discovery by single cell RNA sequencing (scRNAseq) focused on mouse cerebral, visual, and somatosensory cortices (5, 7, 8, 9). More recently, technological advances, including RNAseq using single nuclei (snRNAseq) instead of single cells (10, 11, 12), have extended these investigations into human neuronal cell type discovery (13, 14). Comprehensive reviews of these recent advances have been reported recently (15, 16).

Initial efforts toward human neuronal cell type discovery focused on identifying broad lineages. Pollen *et al.* profiled 65 neuronal cells into six categories - neural progenitor cells, radial glia, newborn neurons, inhibitory interneurons, and maturing neurons (17), while Darmanis *et al.* sequenced 466 cells, also identifying six broad, but distinct, categories - oligodendrocytes, astrocytes, microglia, endothelial cells, oligodendrocyte precursor cells (OPCs), and neurons (18). Darmanis et al. further subtyped the adult neurons into 2 excitatory and 5 inhibitory types. More recent single *nuclei* RNAseq investigations are attempting more comprehensive cell typing. Lake *et al.* sampled 3,227 nuclei from 6 Brodmann areas, from which the neurons were classified into 8 excitatory and 8 inhibitory subtypes (13). Similarly, Boldog *et al.* sampled 769 nuclei from layer 1 of the Middle Temporal Gyrus (MTG) and identified 11 distinct inhibitory cell types (14).

Comparing results between these studies has been challenging given the different areas and layers of cortex sampled. Many of the studies leveraged classical cell type markers derived from the mouse scRNAseq literature. For example, SNAP25 expression was used to broadly define neuronal cells, while GAD1 expression defined inhibitory interneurons. Additional classical markers have then been used to subdivide the excitatory and inhibitory classes, such as CUX2 or VIP respectively; however, these markers individually are still not specific enough to define discrete cell type classes at the level of granularity revealed by clustering of the sc/snRNAseq data. In fact, there has been surprisingly limited overlap in gene sets specific for individual cell type clusters between studies, as the genes found in each study appear to be sensitive to both the context and methodology used. For example, Lake *et al*. found that cluster In1 had CNR1 (Table S5 in reference 13) as the highest ranked marker, while Boldog *et al.* found 7 distinct inhibitory types that expressed this marker (Figure 3 in reference 14). Without a standardized methodology for determining the necessary and sufficient marker genes and a corresponding marker gene reference database, comparison of newly-identified cell types to those reported in previous studies requires a complete reprocessing of the data.

## Application to the human immune system

Single cell transcriptomic analysis has also been applied to study the functional cell type diversity of the human immune system (reviewed in Stubbington 2017 (19)). Bjorklund et al. used scRNAseq to explore the subtype diversity of CD127+ innate lymphoid cells isolated from human tonsil, providing an in-depth transcriptional characterization of the three major subtypes - ILC1, ILC2, and ILC3, and three additional subtypes within the ILC3 class, by comparing their single cell transcriptional profiles (20).

Two recent studies explored the subtype diversity of dendritic cells in human blood. In addition to identifying two conventional dendritic cell subtypes (cDC1 and cDC2) and one plasmacytoid dendritic cell subtype, See et al. identified several subtypes that appear to correspond to precursor cells, including one early uncommitted CD123^+^ pre-DC subset and two CD45RA^+^CD123^lo^ lineage-committed subsets (pre-cDC1 and pre-cDC2), using cell sorting, scRNAseq, and *in vitro* differentiation assays (21). Villani et al. used fluorescence-activated cell sorting and scRNAseq to delineate six different dendritic cell subtypes (DC1 - 6) and four different monocyte subtypes (Mono1 - 4), and showed that these different subtypes, which were defined based on their transcriptional profiles, exhibited different functional capabilities for allogeneic T cell stimulation and for cytokine production following TLR agonist stimulation (22).

Two recent studies have explored the phenotypes of immune cells infiltrating tumor specimens using scRNAseq. In melanoma, Tirosh et al. found that the non-malignant tumor microenvironment was composed of T cell, B cell, NK cell, endothelial cell, macrophage and cancer-associated fibroblast (CAF) subsets (23). In contrast to the distinct transcriptional phenotypes of the malignant component across individual melanoma specimens, common features could be observed in the non-malignant components, with important therapeutic implications. Expression of multiple complement factors by CAFs correlated with the extent of T cell infiltration. T cells with activation-independent exhaustion profiles, characterized by expression of co-inhibitory receptors (e.g. PD1 and TIM3), could be distinguished from cytotoxic T cell profiles. Potential biomarkers that distinguish between exhausted and cytotoxic T cells could aid in selecting patients for immune checkpoint blockade. In hepatocellular carcinoma, Zheng at al. found clonal enrichment of both regulatory T cells and exhausted CD8 T cells using scRNAseq and T cell receptor repertoire analysis (24). The diagnostic and prognostic significance of these findings remain to be explored.

While these studies illustrate the power of single cell genomics in identifying important functional cell subtypes, they also illuminate a major challenge in comparing the results from different studies, due to the lack of a consistent, reusable approach for naming, defining, and comparing new cell types being identified by these high content phenotyping technologies. For example, in the two studies focused on the identification of dendritic cell subtypes, it is unclear if the cDC1 and cDC2 subtypes identified by See et al. correspond to the DC1 and DC2 subtypes identified by Villani et al. Indeed, the only way to make this determination would be to perform a *de novo* comparative analysis of the transcriptional profiles from both studies. For these studies to truly comply with the newly emerging FAIR principles of open data (25), a robust reproducible strategy for defining and representing new cell types will be essential to support their broad interoperability.

## Ontological representation

Biomedical ontologies, as promoted by the Open Biomedical Ontology (OBO) Foundry (26), provide a framework to name and define the types, properties and relationships of entities in the biomedical domain. The Cell Ontology (CL) was established in 2005 to provide a standard reference nomenclature for *in vivo* cell types, including those observed in specific developmental stages in humans and different model organisms (3). The semantic hierarchy of CL is mainly constructed using two core relations - *is_a* and *develops_from.* Masci *et al.* proposed a major revision to the CL using dendritic cells as the driving biological use case in which the expression of specific marker proteins on the cell surface (e.g. receptor proteins) or internally (e.g. transcription factors) would be used as the main *differentia* for the asserted hierarchy (27). Diehl *et al.* applied this approach first to cell types of the hematopoietic system and then later to the full CL (28, 29, 30). As of December 2017, the CL contained 2199 cell type classes, with 583 classes within the hematopoietic cell branch alone.

We recently discussed the challenges faced by the CL in the era of high-throughput, high-content single cell phenotyping technologies, including sc/snRNAseq (4). One of the key recommendations was to establish a standard strategy for defining cell type classes that combine three essential components:

- the minimum set of ***necessary and sufficient marker genes*** selectively expressed by the cell type,
- a ***parent cell class*** in the Cell Ontology, and
- a ***specimen source description*** (anatomic structure + species).

In order to identify the set of necessary and sufficient marker genes from an sc/snRNAseq experiment, we have developed a method - NSforest - that utilizes a random forest of decision trees machine learning approach.

To illustrate how this approach can produce standard cell type definitions, we have applied the method to a transcriptomic dataset derived from single nuclei isolated from the middle temporal gyrus, cortical layer 1 of a post-mortem human brain specimen (Figure 1a in reference 14). Transcriptional profiles obtained from RNA sequencing of a collection of single sorted nuclei was used to identify 16 discrete cell types using an iterative data clustering approach. Based on the expression of the previously charaterized marker genes SNAP25 and GAD1 for broad classes, 11 inhibitory interneurons, 1 excitatory neuron and 4 glial cell type clusters were identified.

**Figure 1.**
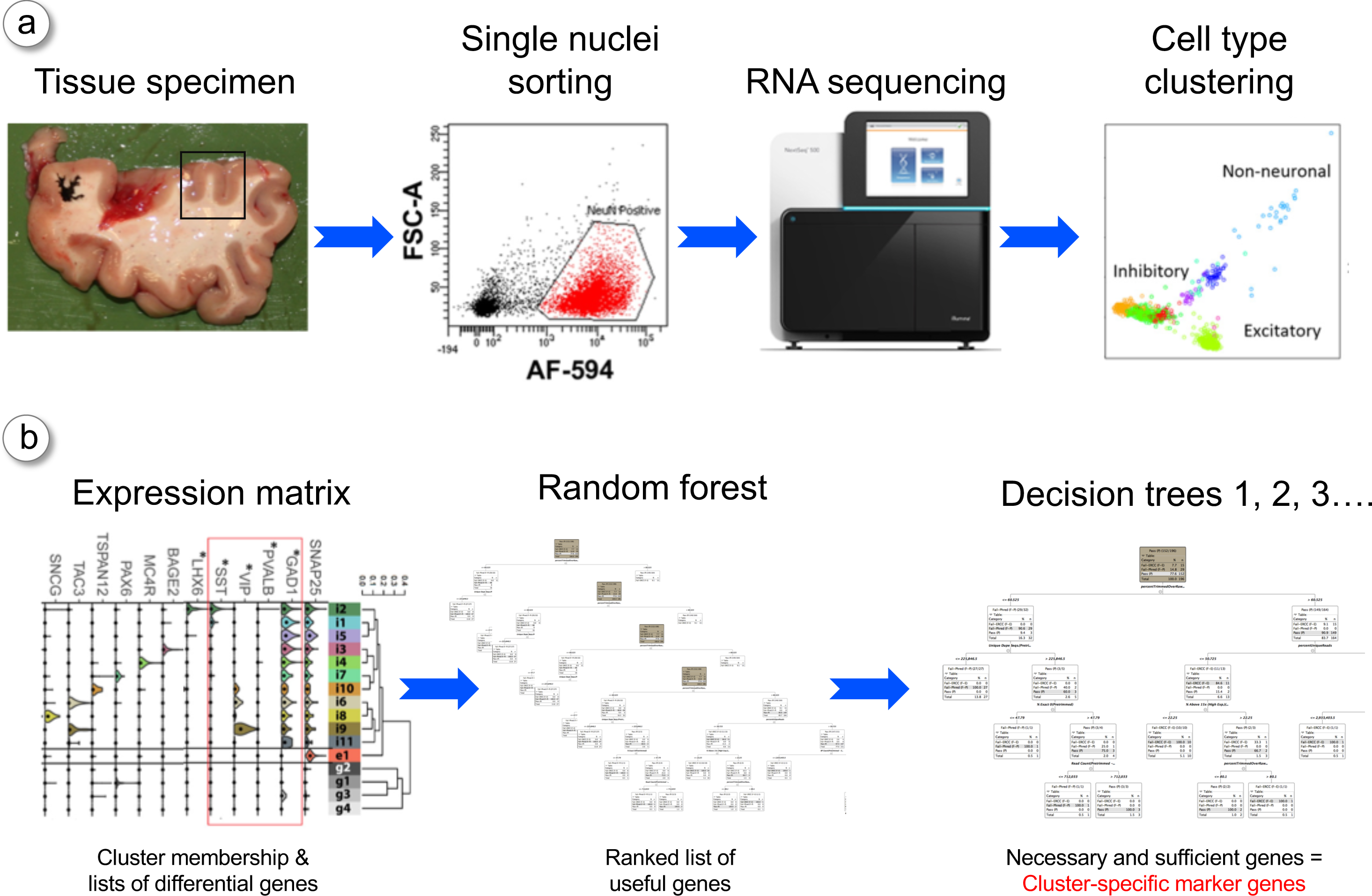
Identification of necessary and sufficient marker genes using NSforest - a) A typical single cell/single nuclei RNA sequencing workflow in which a tissue specimen is obtained, single cells/nuclei isolated by fluorescence activated cell sorting, amplified cDNA quantified by sequencing, and cell types identified by clustering the resultant transcriptional profiles. b) The NSforest approach takes a data matrix of expression values (e.g. transcripts per million reads) of genes (rows) in single cell/nuclei samples (columns) grouped by cell type cluster membership. In the first step, the expression levels of genes are used as features in the random forest machine learning procedure to train classification models comparing single cell/nuclei expression data in one cell type cluster against single cell/nuclei expression data in all other clusters, for every cell type cluster separately, using the Random Forest Learner in KNIME v3.1.2. Each cell type cluster classification model is constructed from one hundred thousand trees using Information Gain Ratio as the splitting criteria, where each decision tree is generated using the default bagging parameters - the square root of the number of features and a bootstrap of samples equal to the training set size. For each cell type cluster classification model, the method outputs usage statistics, including how often each gene is used as a branching criterion and the number of times it was a candidate across all random decision trees. By summing the frequency of use when a candidate across the first three branching levels, the list of genes can be ranked by their usefulness in distinguishing one cell type clusters from the other clusters. In the second step, single decision trees are constructed using the first gene from the ranked list, the first two genes, the first three genes, etc. Each individual tree is then assessed for classification accuracy and tree topology using the training data. Given the objective of determining the necessary and sufficient marker genes, we apply additional criteria in scoring the trees - we restrict each gene to being used in only one branch per tree, and find the optimal classification for the target cluster only, rather than the overall classification score. The addition of genes from the ranked list is stopped when an optimal classification or stable tree topology is achieved. The minimum number of genes used to produce this optimal result corresponds to the set of necessary and sufficient marker genes required to define the cell type cluster.

In the first step (Figure 1b), NSforest takes the gene expression data matrix of single nuclei with their cell type cluster membership as input, and develops a classification model for each cell type cluster by comparing each Cluster X versus all non-Cluster X profiles using the random forest algorithm (31). In addition to the classification model itself, NSforest produces a ranked list of features (genes) that are most informative for distinguishing between Cluster X and all of the other clusters.

In the second step, NSforest constructs single decision trees using first the top gene, then the top two genes, top three genes, etc., until a stable tree topology and optimal classification accuracy is achieved. The minimum number of genes necessary to obtain this stable classification result corresponds to the necessary and sufficient set of marker genes defining each cell type cluster within this experimental context.

The expression of the complete set of marker genes obtained from applying NSforest to the single nuclei dataset is illustrated in Figure 2. In most cases, the expression of three marker genes is sufficient to define a cell type cluster, with a range of one to five necessary and sufficient marker genes per cluster. Glial cell subtypes appear to be more distinct from each other, requiring relatively few genes to sufficiently define the cell type. In contrast, neuronal subtypes appear to be more similar, requiring more genes to achieve specificity. In some cases, a combination of both positive and negative expression optimally defines a cell type cluster.

**Figure 2.**
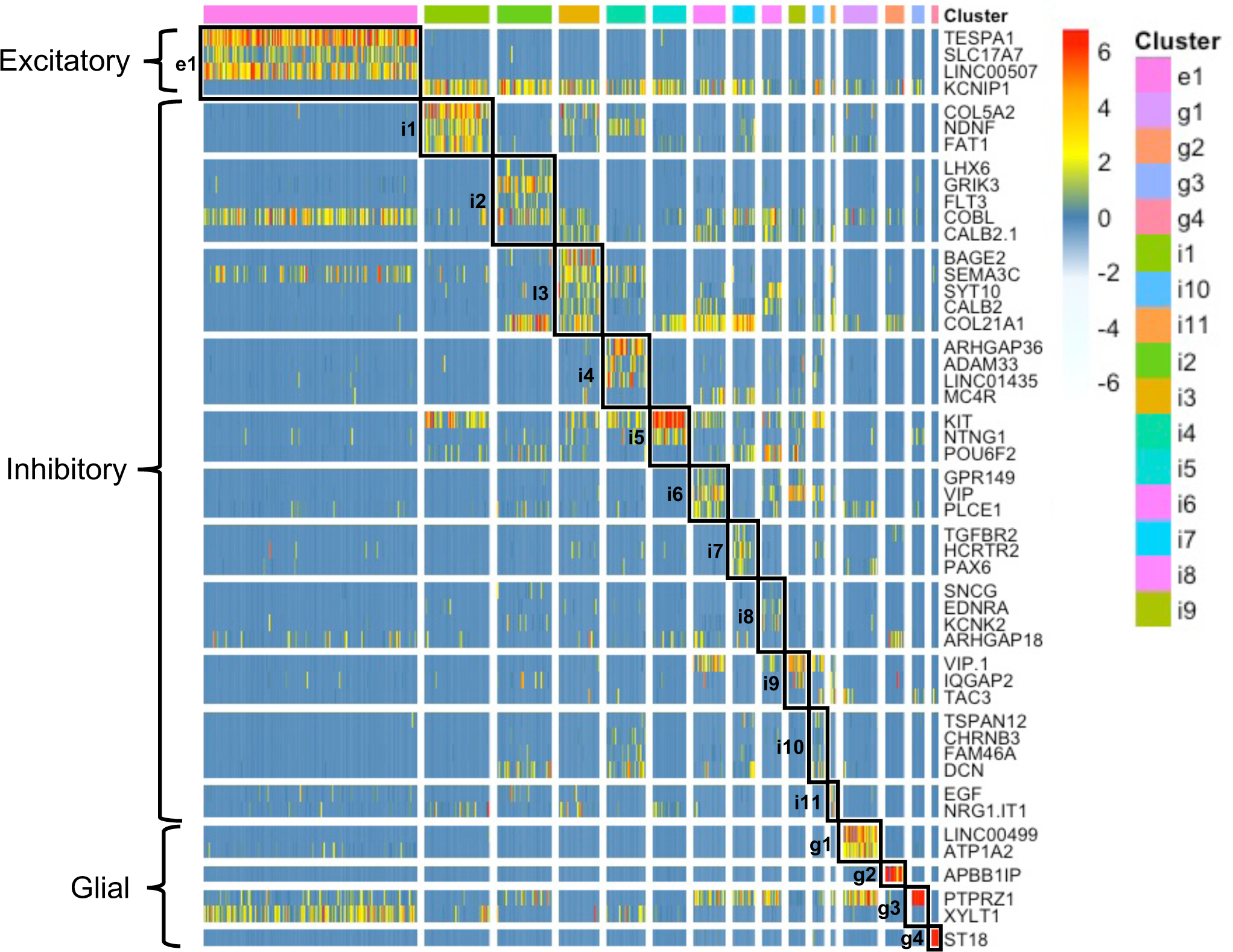
Marker gene expression patterns in single nuclei grouped by cluster - A heatmap of expression levels for the necessary and sufficient marker genes identified for all 16 clusters across all single nuclei grouped by cell type cluster is shown, including 1 excitatory (e1), 11 inhibitory (i1 - i11), and 4 glial (g1 - g4) cell type clusters. In total, 49 markers genes were selected as being necessary and sufficient to distinguish these 16 different cell type clusters from cortical layer 1 of the human brain middle temporal gyrus region.

For one of the inhibitory interneuron cell types defined in this study (i5), we were able to connect the distinct transcriptional profile with a previous cell type defined based on its unique cellular morphology - the Rosehip cell (14). This then allows us to construct an ontological representation that includes both a colloquial name, an alternative name, and a definition combining the necessary and sufficient marker genes, a CL parent cell class, and specimen source information, as follows:

- Colloquial name - ***rosehip neuron***
- Alternative name - ***KIT-expressing MTG cortical layer 1 GABAergic interneuron, human***
- Definition - ***A human middle temporal gyrus cortical layer 1 GABAergic interneuron that selectively expresses KIT, NTNG1, and POU6F2 mRNAs***

A complete set of cell type names and definitions for all cell type clusters identified in this experiment is provided in Table 1.

**Table 1.**
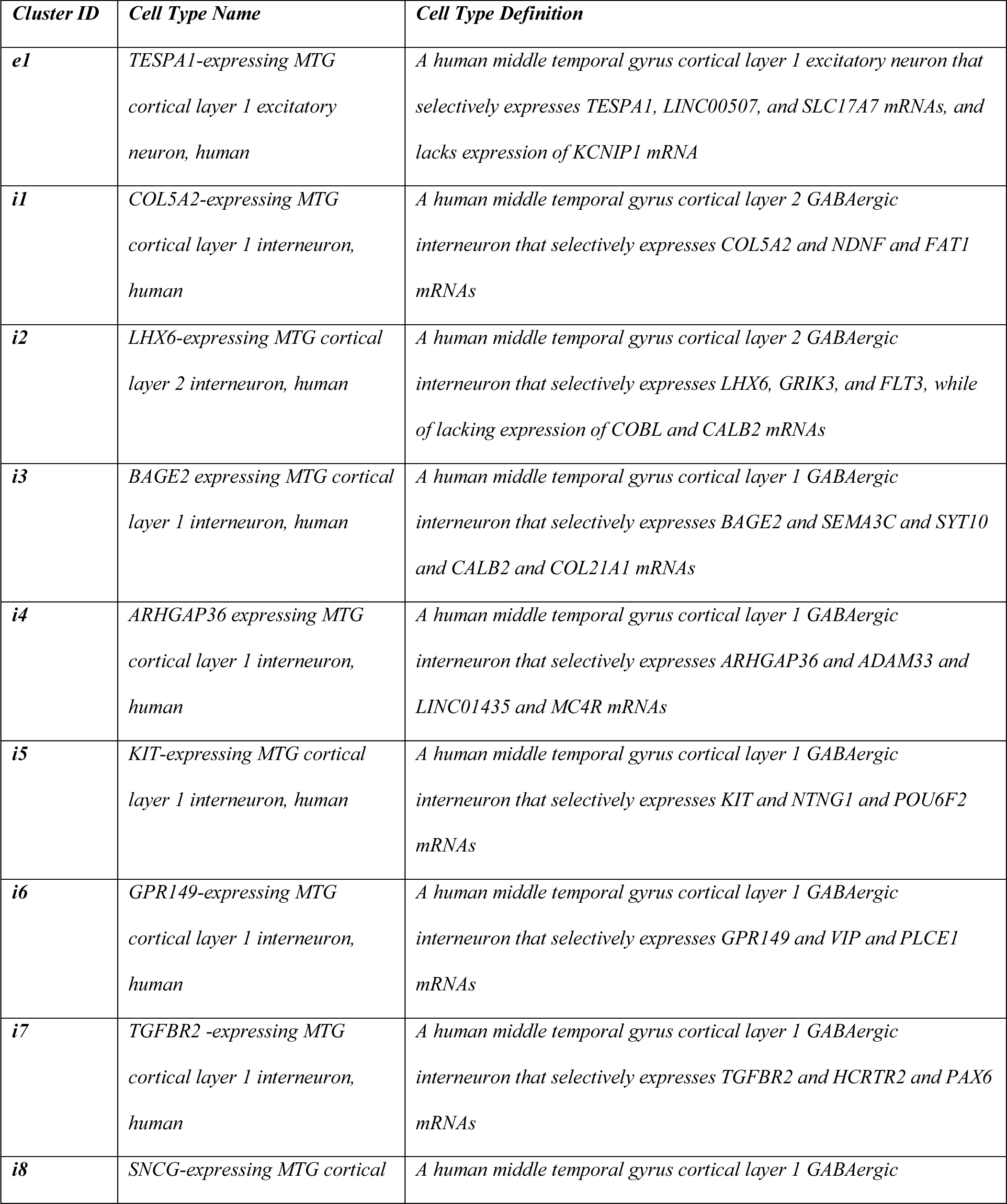
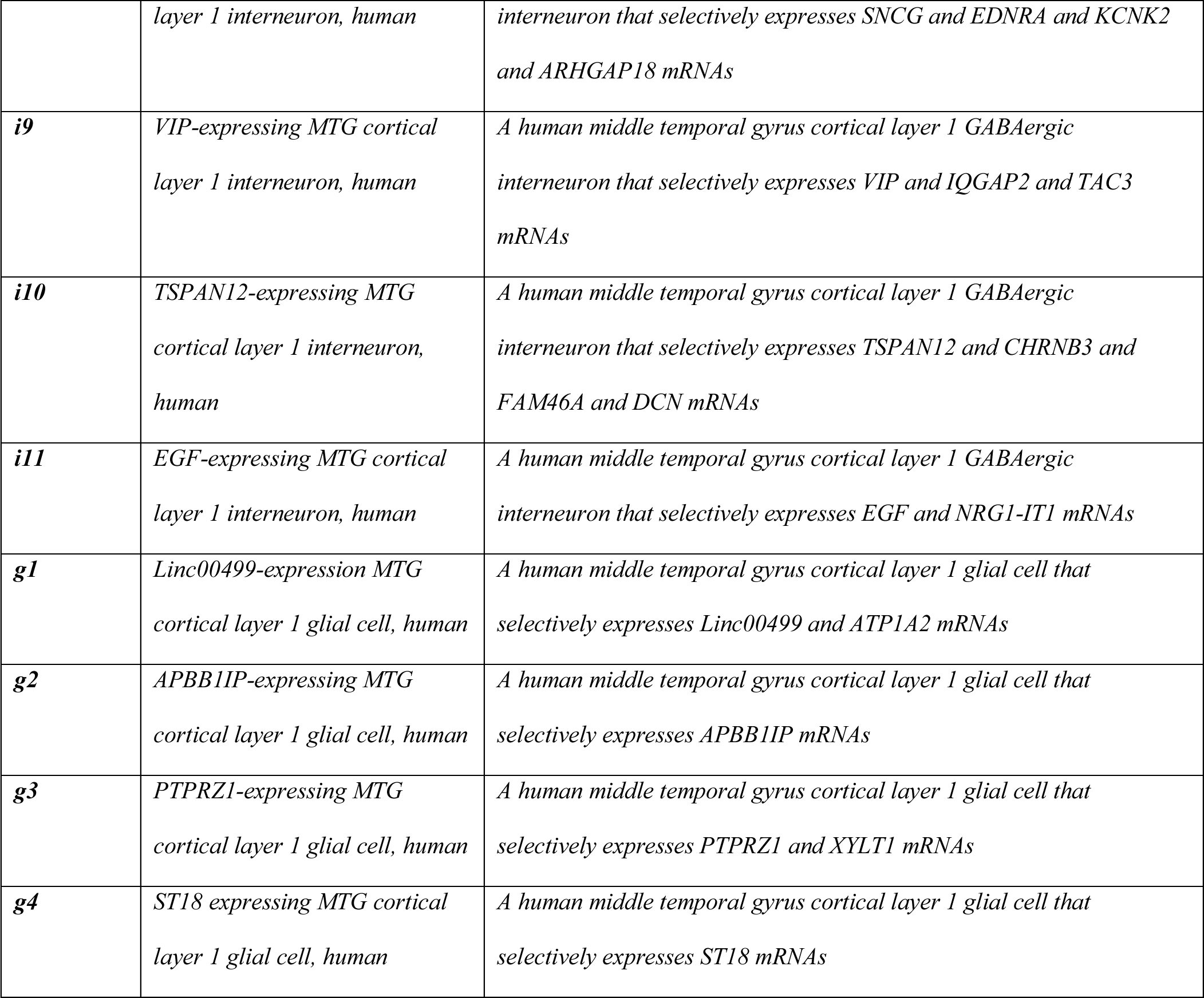
Cell types identified in cortical layer 1 of the human middle temporal gyrus

These informal textual definitions can then be converted into formal ontological definitions, represented in OWL as equivalent classes, using a set of logical axioms that combine assertions about the parent cell class (interneuron), anatomic locations of the neuron cell body (soma), functional capacity of the cell type (gamma-aminobutyric acid secretion), and marker gene expression (expresses some KIT) requirements (Figure 3). Using semantic reasoners, these logical axioms can then be used to infer novel characteristics, e.g. subClass Of ‘cerebral cortex GABAergic interneuron’.

**Figure 3.**
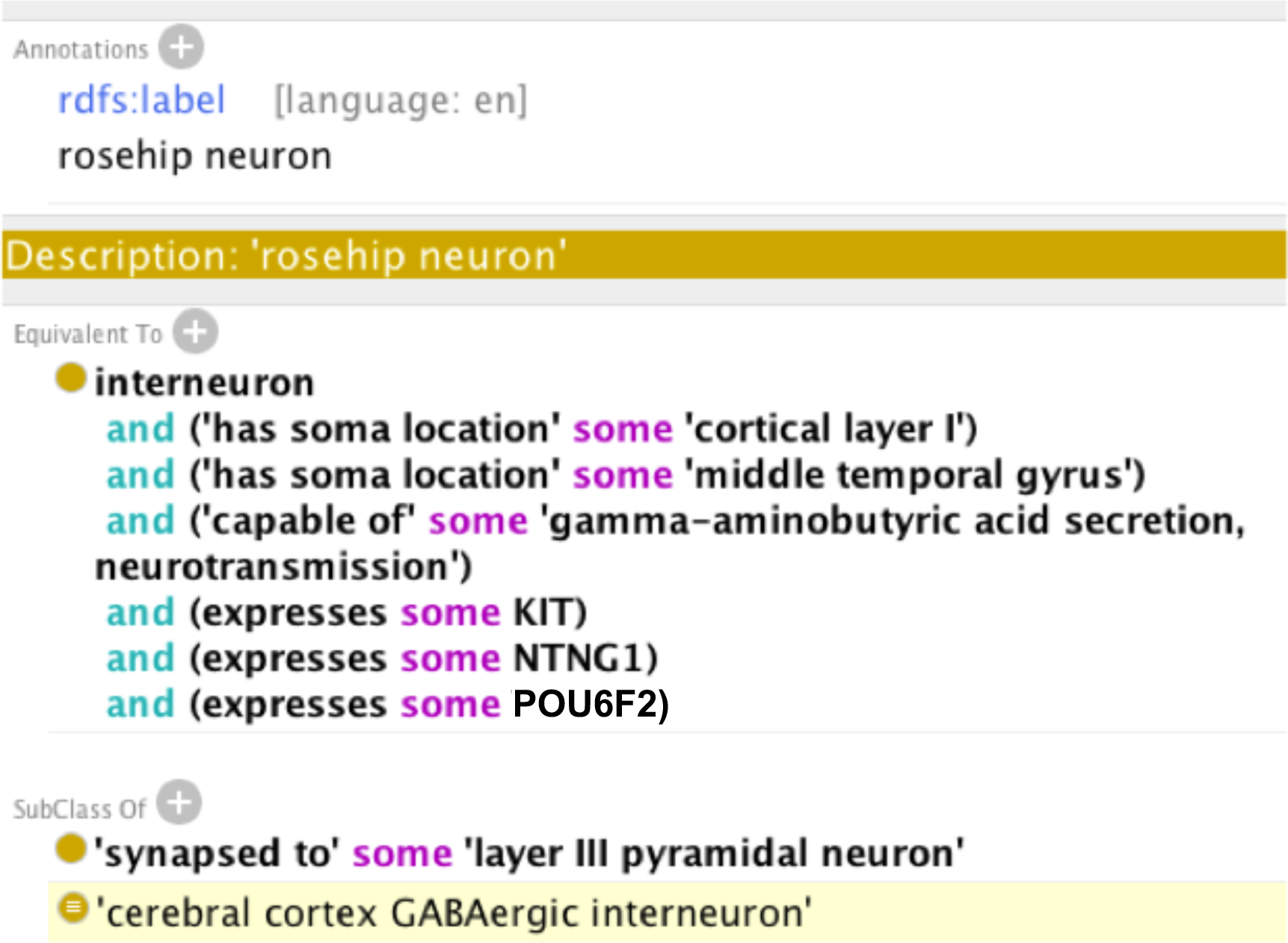
Formal rosehip neuron definition using logical axioms - A set of logical axioms about the anatomic local of the cell body (soma), the functional capacity, and the necessary and sufficient marker gene expressions are combined to construct an equivalent class cell type definition for the rosehip neuron interneuron cluster - i4 (see Boldog 2017 (14) for more information about how this cell type was characterized).

The challenge remains of ensuring that these cell type definitions, whose necessary and sufficient conditions are derived from analysis of data from one particular methodology (scRNAseq), are compatible with both existing cell type classes in the CL and cell types defined using alternative experimental methods and data analysis approaches. Working with CL developers, we are now establishing an extension ontology module containing provisional definitions for novel cell types that we and other research groups will contribute. Ontological reasoners will be used to link these cell types to more general classes in the CL proper, structure them into an extended hierarchy, and determine when separate research groups have defined similar or identical cell types. CL developers will review these provisional cell types periodically to determine when multiple lines of evidence provide sufficient support to promote particular cell type classes to the CL proper. In this way we will ensure the integrity of the CL reference, while still allowing for the rapid expansion of its content to accommodate cell types defined via these new technologies.

## Conclusions

The application of high-throughput/high-content cytometry and single cell genomic techniques is producing an explosion in the number of distinct cellular phenotypes being identified in human specimens. For biomedical ontologies to stay relevant, it will be critical for ontology developers to establish procedures for the processing and incorporation of representations derived from these data-intensive technologies into reference ontologies in a timely fashion. The representation of defined cell types and their relationships in the CL will serve as a reference knowledgebase to support interoperability of information about the role of cellular phenotypes in human health and disease.

## Acknowledgements

This work was supported by the Allen Institute for Brain Science, the JCVI Innovation Fund, the U.S. National Institutes of Health R21-AI122100 and U19-AI118626, and the California Institute for Regenerative Medicine GC1R-06673-B. We thank Nik Schork, Jamison McCorrison, Pratap Venepally, Lindsay Cowell, Bjoern Peters, and Sirarat Sarntivijai for helpful discussion.

